# Exploring the prognostic value of SCARF1 in non-small cell lung cancers

**DOI:** 10.1101/2021.01.28.428617

**Authors:** Daniel A. Patten, Shishir Shetty

## Abstract

Scavenger receptor class F member 1 (SCARF1) has previously been shown to be highly expressed within the human liver, hold prognostic value in hepatocellular carcinoma and mediate the specific recruitment of leukocytes to liver sinusoidal endothelial cells; however, to date, the liver remains the only major organ in which SCARF1 has been explored in any detail. Here, we utilised publically-available RNA-sequencing data from The Cancer Genome Atlas (TGCA) to identify the lungs as a site of significant *SCARF1* expression and attribute the majority of its expression to endothelial cell populations. Next, we show that *SCARF1* expression is significantly reduced in two histologically distinct types of non-small cell lung carcinoma cancers (NSCLCs), lung adenocarcinoma (LUAD) and lung squamous cell carcinoma (LUSC), compared to non-tumoural tissues. Interestingly, loss of *SCARF1* expression was associated with aggressive tumour biology in LUAD tissues, but not in LUSC. Furthermore, increased *SCARF1* expression was highly prognostic of better overall survival in LUAD tumour tissues, but this was again in contrast to LUSC tumours, in which *SCARF1* held no prognostic value. Finally, we showed that *SCARF1* is widely expressed in tumour endothelial cells of non-small cell lung cancers and that its total expression in LUAD tumour tissues correlated with immune score and CD4^+^ T cell infiltration. This study represents the first detailed exploration of *SCARF1* expression in normal and diseased human lung tissues and further highlights the prognostic value and therapeutic potential of SCARF1 in immunologically active cancers.

## Introduction

Globally, lung cancer is the leading cause of cancer mortality and is responsible for more than 1.7 million deaths worldwide each year [1]. This is largely due to the fact that the 5-year survival from lung cancer ranges from 4-17 %, depending on cancer stage and regional variation [2]. Non-small cell lung cancers (NSCLCs) make up the majority (~85 %) of lung cancers [3]; of those, lung adenocarcinoma (LUAD) is the most common histologic subtype, accounting for about 40% of total lung cancer incidence [4]. The second most prevalent subtype of NSCLC is lung squamous cell carcinoma (LUSC), which accounts for around 30% of the global lung cancer incidence [4]. The high mortality rate in lung cancers is largely due to the fact that many patients present with advanced disease at diagnosis, which often includes the presence of metastatic disease [3]. In early stage disease, surgical resection remains the single most effective treatment [2]; in cases where surgery is not a viable option, for example in later stage diseases, combination therapies of thoracic radiotherapy and non-specific chemotherapies have traditionally been used [2]. However, lung cancers are notoriously heterogeneous diseases [5,6] and understanding their molecular make-up is crucial for the development of more targeted therapies. Over the last 20+ years, genetic testing has resulted in a number of highly specific therapies being developed and survival time of patients have improved [2]; nevertheless, furthering our understanding of the tumour immune microenvironment is still pertinent to the development of novel biomarkers and therapies in NSCLCs. Much of the research to date has predominantly focused on immune cell profiles [7,8] and stromal (e.g. fibroblast) populations [9,10] of NSCLC tumours, whilst other tumour-resident cells, such as endothelial cells, are less well studied. We have previously described the prognostic value of the endothelial-expressed scavenger receptor class F, member 1 (SCARF1) in hepatocellular carcinoma (HCC) and hypothesised that it plays a key role in the recruitment of proinflammatory CD4^+^ T cells to the HCC tumour microenvironment [11]; here, we sought to study SCARF1 in the context of NSCLCs.

Scavenger receptors are a large super-family of proteins which are defined by their ability to bind and internalise a vast range of endogenous and exogenous ligands and removing them from the general circulatory system [12]. Consequently, scavenger receptors represent a major subset of innate pattern recognition receptors (PRRs) and are known to key roles in tissue homeostasis, infections and inflammatory diseases [12]. Given the high environmental antigenic load that is inhaled with each breath, it is unsurprising that a number of scavenger receptors are expressed in resident cell populations of the lung. In particular, the Class A family of scavenger receptors is known to play a key role in innate immunity in response to a number of microbial infections [13,14], environmental nanoparticles and allergens [15,16] and inhaled oxidants [17]. SCARF1 has also previously been confirmed to be expressed in human [18] and murine [19,20] lung tissues. SCARF1 was first identified in cDNA libraries from human umbilical vein endothelial cells (HUVEC) [21] and exhibits high expression in primary human liver sinusoidal endothelial cells (LSEC) [22], which are also known to be exposed to high levels of antigenic matter from the gut. SCARF1 has been shown to bind and internalise a wide range of endogenous and exogenous ligands [23], such as apoptotic host cells [24] and viral [25–27], fungal [19] and bacterial [28–31] antigens. Nevertheless, its expression in human lung tissues and in cancers of the lung have not been explored in any detail to date.

Here, through the utilisation of the publically-available TGCA (The Cancer Genome Atlas) datasets (http://cancergenome.nih.gov), we initially described the expression of *SCARF1* in a range of human tissues and demonstrate lung tissues as a major site of expression. Next, we showed that the expression of *SCARF1* within lung tissues is largely associated with endothelial cell populations. Subsequently, we described a downregulation of *SCARF1* expression in two distinct histological types of non-small cell lung carcinoma tumours, lung adenocarcinoma (LUAD) and lung squamous cell carcinoma (LUSC), compared to non-tumourous control tissues. Following this, we explored the relationship of *SCARF1* expression with tumour progression and, consequently, found an association with loss of *SCARF1* expression with aggressive tumour biology in LUAD tumours, but not in LUSC tumour tissues. Following this, we evaluated the prognostic value of *SCARF1* expression in LUAD and LUSC tumours by generating survival curve data, via KM Plotter (http://kmplot.com/analysis/). In support of the pathological findings, high *SCARF1* expression in LUAD tumour tissues was found to correlate with a better overall survival; however, *SCARF1* expression showed no prognostic value in LUSC tumours. In addition, through use of the LungECtax database (https://endotheliomics.shinyapps.io/lungectax/), we demonstrated *SCARF1* expression across a range of of endothelial cell populations present in both normal lung and NSCLC tumour tissues. Finally, we used publically-available tools, Estimation of STromal and Immune cells in MAlignant Tumor tissues using Expression data (ESTIMATE; https://bioinformatics.mdanderson.org/estimate/) and Tumor IMmune Estimation Resource (TIMER; https://cistrome.shinyapps.io/timer/) to correlate *SCARF1* expression with immune score and the level of CD4^+^ T cell infiltration, respectively, in LUAD tumour tissues. Using ESTIMATE, we demonstrated a moderate positive correlation between *SCARF1* expression and immune infiltration score and, via TIMER, we confirmed that *SCARF1* expression correlates more specifically with CD4^+^ T cell infiltration. Our results demonstrate that *SCARF1* could be a prognostic biomarker in LUAD and that lung-expressed SCARF1 could potentially function to alter the inflammatory status of the tumour microenvironment.

## Materials and Methods

### In silico *data analysis*

Publically-available RNA-sequencing data was utilised throughout this study and several publically-available tools were used in its analysis. To explore *SCARF1* expression in range of human major organs and in lung tumour and relevant non-tumourous tissue controls, data was obtained from the Genotype-Tissue Expression (GTEx) and The Cancer Genome Atlas (TGCA) datasets, via the University of California Santa Cruz (UCSC) Xena tool (https://xenabrowser.net/). *SCARF1* expression data in various cell populations of the human lung was generated by the LungMAP Consortium and downloaded from (www.lungmap.net) (accessed on 4th June 2020). Correlation of *SCARF1* expression with tumour progression/aggression was performed via the cBioPortal website (https://www.cbioportal.org/) (accessed 28^th^ April 2020). With the use of the publically-accessible tool KM Plotter (http://kmplot.com/analysis/), survival data was generated using the Affymetrix ID 206995_x_at (*SCARF1*). Data was censored at a 60-month threshold and was split into two groups (‘High’ and ‘Low’) by the median of *SCARF1* expression. Resultant data was exported to Prism^®^ 6 software (GraphPad Software Inc.) and survival curves were produced. tSNE plots of SCARF1 expression in normal endothelial cells (NEC) and tumour endothelial cells (TEC) were generated by the publically-available LungECtax database (https://endotheliomics.shinyapps.io/lungectax/; accessed on 4^th^ June 2020). Immune infiltration scores of LUAD tumours were correlated with *SCARF1* expression via Estimation of STromal and Immune cells in MAlignant Tumor tissues using Expression data (ESTIMATE; https://bioinformatics.mdanderson.org/estimate/; accessed 19^th^ August 2020). Level of CD4^+^ T cell infiltration of LUAD tumours was correlated with *SCARF1* expression via the Tumor IMmune Estimation Resource (TIMER; https://cistrome.shinyapps.io/timer/; accessed 12th May 2020).

### Statistical analyses

All data were tested for normal distribution by the D’Agostino-Pearson omnibus test. All data were found to be non-parametric and so were expressed as median ± interquartile range (IQR), with the number of experimental repeats (*n*) specified in each case. For single comparisons, statistical significance was determined by Mann-Whitney *L·*-test, whereas evaluation of multiple treatments was performed by Kruskall-Wallis one-way analysis of variance with post hoc Dunn’s test. Matched data was analysed by a Wilcoxon signed-rank test. A *p*-value of ≤ 0.05 was considered as statistically significant. All statistical analyses were undertaken using Prism^®^ 6 software (GraphPad Software Inc.).

## Results

### SCARF1 *is highly expressed in human lung tissues*

We have previously shown that SCARF1 is highly present in in human liver tissues [11,22]; however, the exploration of its expression in other major organs is limited. Here, through the analysis of the publically-available TCGA RNA sequencing data and using the expression in the liver as a reference, we showed that *SCARF1* was differentially expressed at the gene level in a wide range of human major organs. In the 14 major organs explored, 8 (pancreas, brain, skin, bladder, kidney, esophagus, stomach and colon) demonstrated significantly lower expression than the liver (Figure 1A), whereas 4 (heart, breast, lung and spleen) exhibited significantly higher expression (Figure 1A); the only exception was the small intestine, which showed comparable *SCARF1* expression to the liver. Of the tissues explored, the lungs exhibited marked *SCARF1* expression, with only the levels in the spleen found to be higher. Through the use of additional publically-available RNA sequencing data generated from bulk cell populations, we show that endothelial cell populations account for the vast majority of *SCARF1* expression in the human lung at all stages of the life cycle (neonate-infant-child-adult), when compared with epithelial, mesenchymal and immune cell populations (Figure 1B).

**Figure 1.**
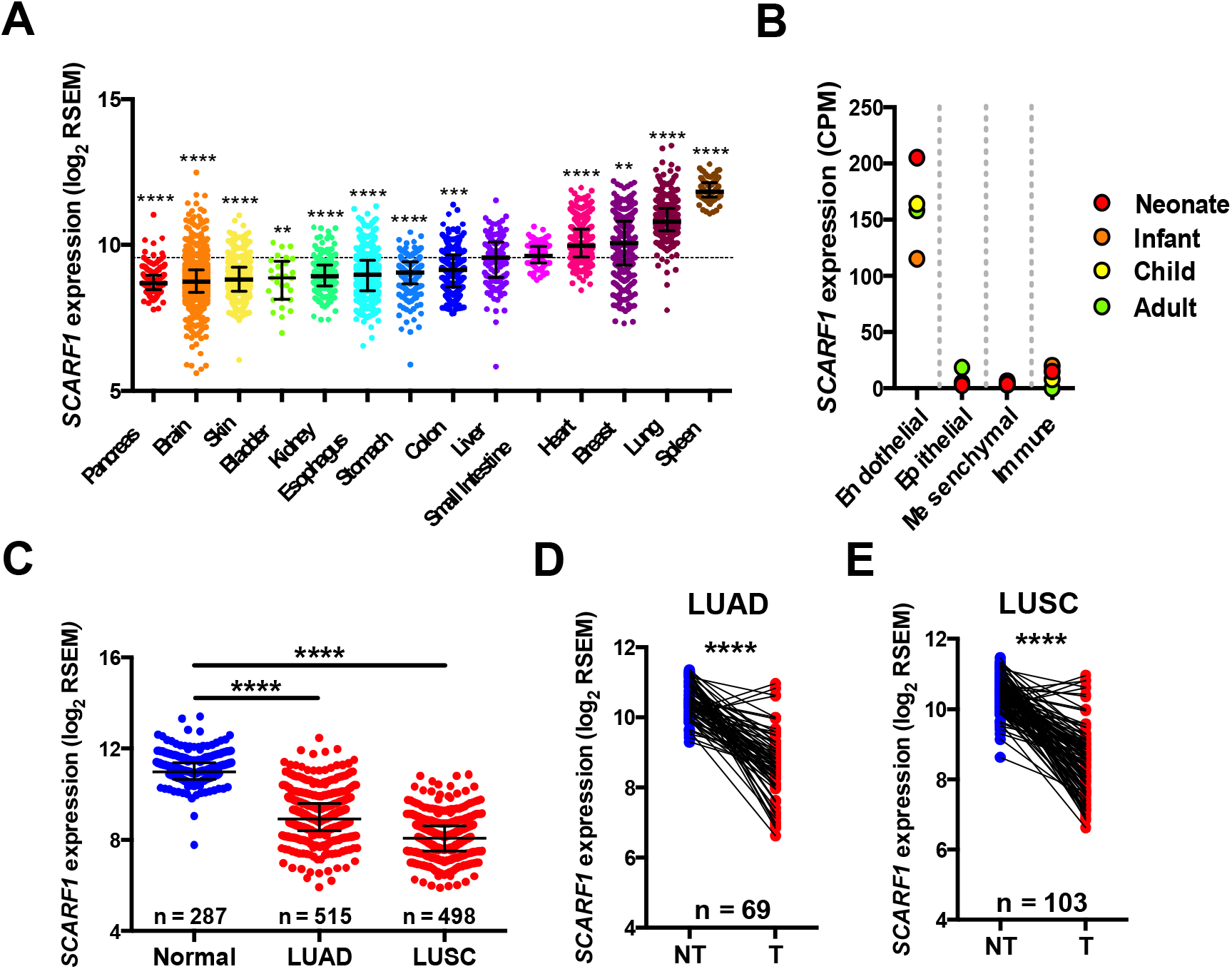
SCARF1 mRNA expression is high in lung tissues, but downregulated in lung cancers. (A) Expression of SCARF1 gene expression in major tissues: pancreas (n = 169); brain (n = 1136); skin (n= 556); bladder (n = 28); kidney (n = 167); esophagus (n = 662); stomach (n = 208); colon (n = 345); liver (n = 160); small intestine (n = 92); heart (n = 376); breast (n = 292); lung (n = 396); spleen (n = 99). *, **, *** and **** are representative of statistical significance as measured by the Kruskal-Wallis test, where *p*□≤□0.05, *p*□≤□0.01, *p*□≤□0.005 and *p*□≤□0.001, respectively. RSEM = RNA-Seq by Expectation Maximization. The dotted line is representative of the *SCARF1* expression in Liver. (B) SCARF1 gene expression in different bulk cell populations of neonatal (red), infant (orange), child (yellow) and adult (green) human lung. CPM = counts per million. (C) Comparison of SCARF1 gene expression in normal lung tissues with lung adenocarcinoma (LUAD) and lung squamous cell carcinoma (LUSC) tumour tissues. **** indicates statistical significance as measured by the Kruskal-Wallis test, where p□≤□0.001. (D and E) Comparison of SCARF1 gene expression in non-tumoural (NT) tissues with matched tumoural (T) tissues in both LUAD and LUSC. **** indicates statistical significance as measured by a Wilcoxon signed-rank test, where p□≤□0.001. Data in (A) and (C-E) was generated from the TGCA dataset using the University of California Santa Cruz (UCSC) Xena tool (https://xenabrowser.net/). Data in (B) are based upon data generated by the LungMAP Consortium and downloaded from (www.lungmap.net), on 4^th^ June 2020. The LungMAP consortium and the LungMAP Data Coordinating Center (1U01HL122638) are funded by the National Heart, Lung, and Blood Institute (NHLBI).

### *SCARF1* expression is downregulated in NSCLC

We have previously shown that SCARF1 is downregulated in gastrointestinal cancers, such as hepatocellular carcinoma [11], and, given the high expression in the human lung, we next explored its expression in non-small cell lung carcinomas (NSCLCs). Analysis of the publically-available RNA-sequencing data from The Cancer Genome Atlas (TGCA) and Genotype-Tissue Expression (GTEx) project showed that *SCARF1* expression is significantly (*p*□≤□0.001) lower in both LUAD and LUSC tumour tissues in comparison to normal lung tissues (Figure 1C). The downregulation of *SCARF1* expression was also evident when we compared tumour tissues with matched non-tumourous tissues from the same donor in both LUAD (Figure 1D) and LUSC (Figure 1E).

### *Loss of* SCARF1 *expression is associated with more advanced and aggressive tumours in LUAD, but not LUSC*

Previously, we have shown that a loss of SCARF1 expression in hepatocellular carcinoma tumours is associated with more aggressive tumour biology [11]; here, we aimed to utilise the TGCA dataset to explore whether this was also the case in NSCLCs. Firstly, we explored *SCARF1* expression levels in cases of NSCLCs at different stages of the disease, from early stage disease (Stage I) through to highly developed and metastatic disease (Stage IV). When compared to patients with Stage I disease, cohorts of LUAD patients with Stages II, III and IV disease all demonstrated a trend for decreased *SCARF1* expression; however, only the data for the Stage II cohort was calculated to be statistically significant (*p*□≤□0.005). We next correlated *SCARF1* expression with other parameters commonly associated with tumour aggressiveness; in particular, we focussed on Aneuploidy Score [33] and Buffa Hypoxia Score [34]. We demonstrated a moderate negative correlation of *SCARF1* expression with both Aneuploidy Score (Figure 2B) and Buffa Hypoxia Score (Figure 2C) in LUAD tumour tissues, thus providing further evidence that a loss of *SCARF1* expression is associated with more adverse tumour biology. In stark contrast, patient cohorts with LUSC all showed comparable *SCARF1* expression levels, regardless of disease stage (Figure 2D). In addition, the negative correlations between *SCARF1* expression levels and Aneuploidy Score and Buffa Hypoxia Score were much weaker in LUSC tumour tissues (Figure 2E and 2F), when compared to those seen in LUAD tumours (Figures 2B and 2C).

**Figure 2.**
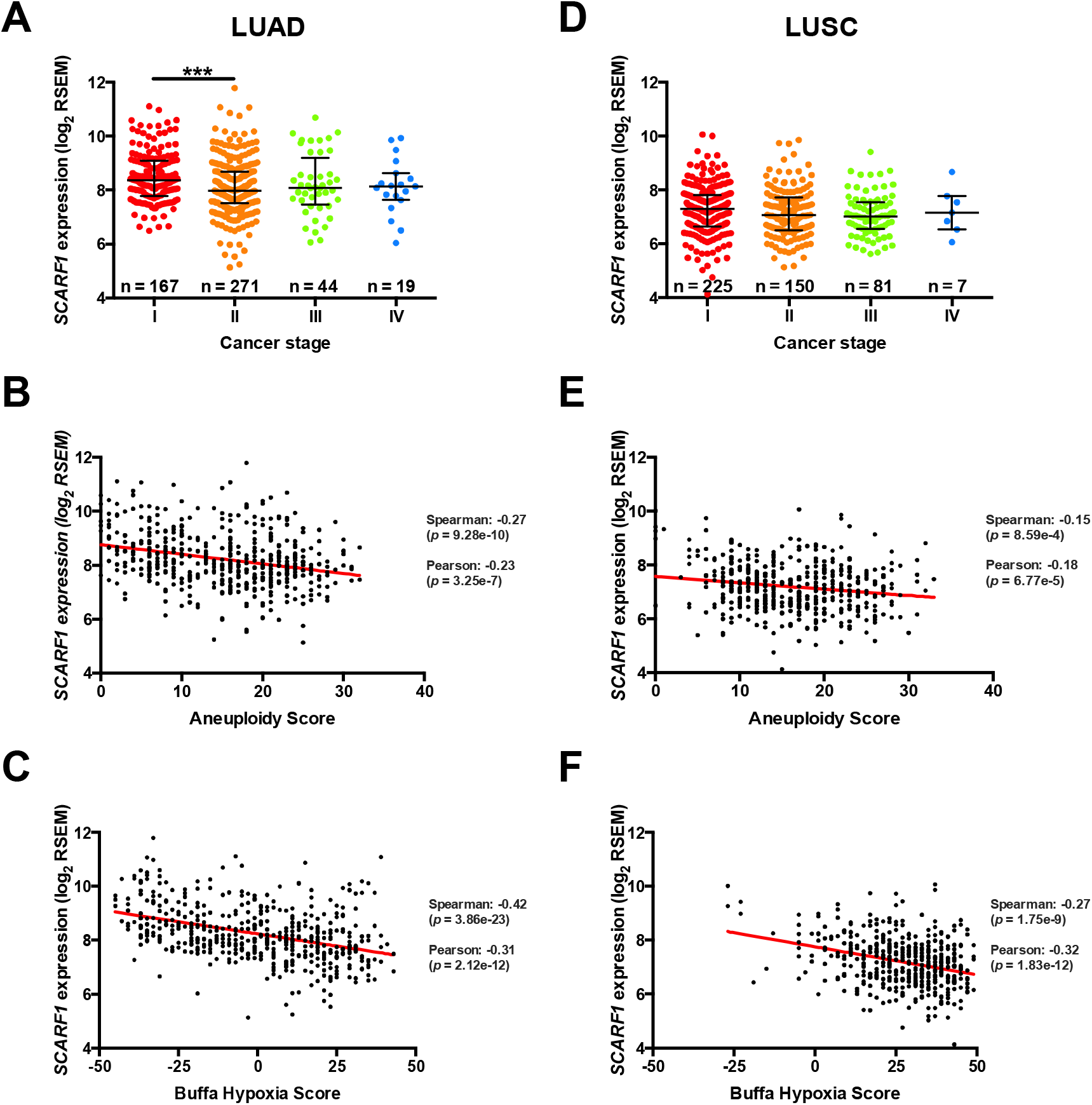
More advanced and aggressive LUAD tumours exhibit lower *SCARF1* expression. (A) *SCARF1* expression in LUAD tumour tissues from the four cancer stages. *** is representative of statistical significance as measured by the Mann Whitney *U*-test, where p□≤□0.005. *SCARF1* expression in LUAD tumours correlated to tumour aggression parameters (B) Aneuploidy score (n = 493) and (C) Buffa Hypoxia score (n = 503). (D) *SCARF1* expression in LUSC tumour tissues from the four cancer stages. *SCARF1* expression in LUSC tumours correlated to tumour aggression parameters (E) Aneuploidy score (n = 465) and (F) Buffa Hypoxia score (n = 466). Data in this Figure was generated from the TGCA dataset using the cBioPortal website (https://www.cbioportal.org/) (accessed 28^th^ April 2020).

### *Loss of* SCARF1 *expression is associated with worse outcomes in LUAD, but not LUSC*

We next aimed to explore if *SCARF1* expression was in any way associated with outcome in either histological subtype of NSCLC. Firstly, we explored disease-free status since initial treatment in LUAD and showed that patients whose disease recurred or progressed had significantly (*p*□≤□0.05) lower expression levels of intratumoutal *SCARF1* expression than those who remained disease-free (Figure 3A). Following this, we explored *SCARF1* expression and disease-specific survival status in LUAD patients; *SCARF1* expression in tissues from those patients who subsequently died with a tumour burden was significantly (*p*□≤□0.005) lower than patients who remained alive or died tumour-free (Figure 3B). We next explored the same parameters in LUSC patients. We demonstrated comparable levels of *SCARF1* between patients whose disease recurred/progressed and those who remained disease-free (Figure 3C) and between those patients who subsequently died with a tumour burden and those who remained alive or died tumour-free (Figure 3D).

**Figure 3.**
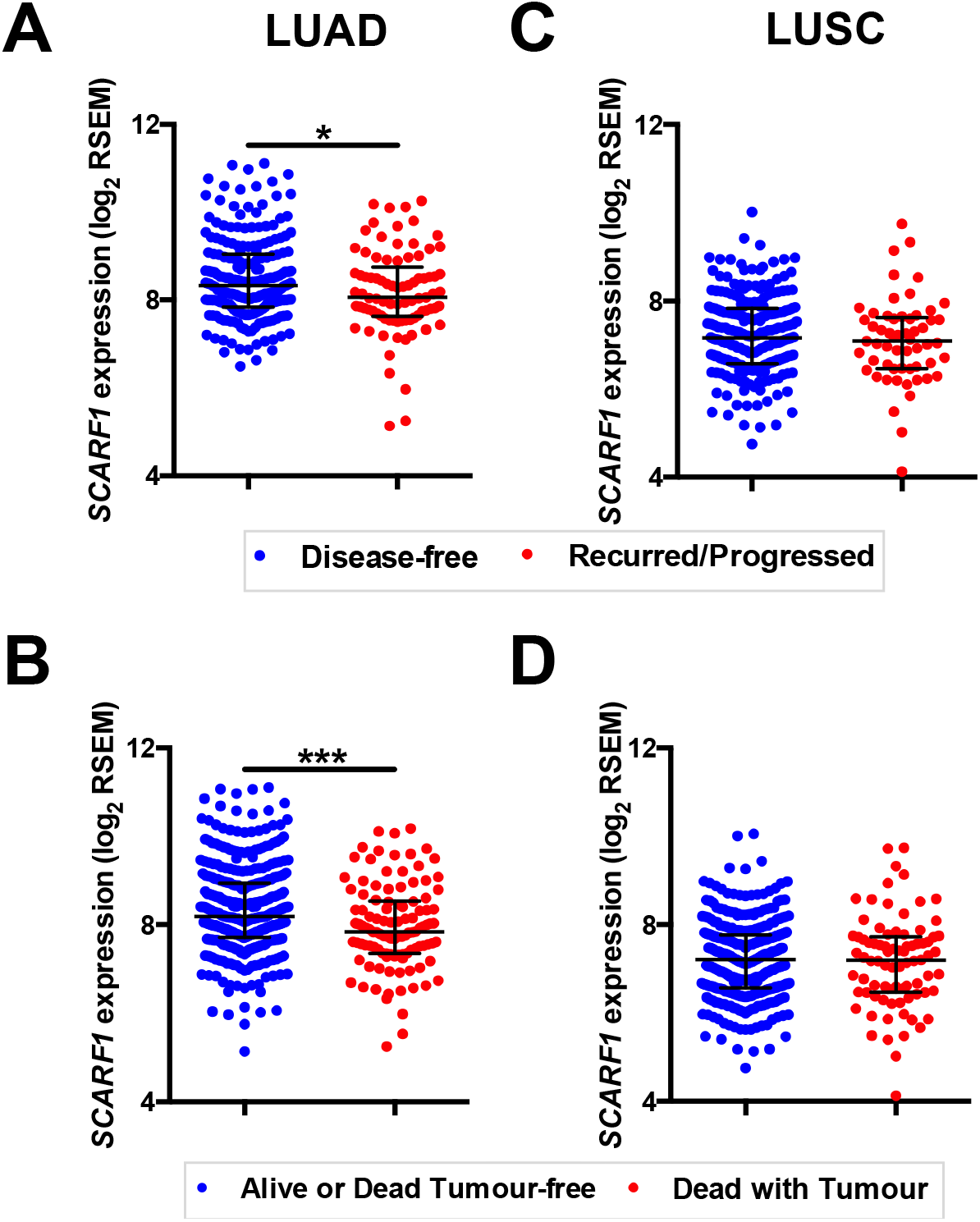
Lower *SCARF1* expression is associated with worse outcomes in LUAD, but not LUSC. Comparison of *SCARF1* expression in (A) LUAD and (C) LUSC patients who survived and remained disease-free (blue dots) and those in which disease recurred or progressed (red dots). * is representative of statistical significance as measured by the Mann Whitney U-test, where p□≤□0.05. Comparison of *SCARF1* expression in (B) LUAD and (D) LUSC patients who survived or died tumour-free (blue dots) and those who died with tumour present (red dots). *** is representative of statistical significance as measured by the Mann Whitney U-test, where p 0.005. Data in this Figure was generated from the TGCA dataset using the cBioPortal website (https://www.cbioportal.org/) (accessed 28^th^ April 2020).

### *Prognostic value of* SCARF1 *expression in NSCLCs*

Having found that a loss of *SCARF1* expression correlates with more advanced and aggressive tumours (Figure 2) and worse outcome (Figure 3) in LUAD, we next sought to investigate its prognostic value in NSCLCs. With regards to overall survival, high expression of *SCARF1* was highly indicative of a better prognosis in LUAD (HR = 0.68, 95 % CI = 0.52 – 0.87, *p* ≤ 0.01; Figure 4A). We also assessed the prognostic value of *SCARF1* expression in correlation with a range of clinicopathological features in LUAD. In both male and female patients, higher *SCARF1* expression was suggestive of better overall survival (*Male* HR = 0.59, 95 % CI = 0.41 – 0.85, *p* ≤ 0.005; *Female* HR = 0.61, 95 % CI = 0.40 – 0.94, *p* ≤ 0.05; Figure 4B) 0.61 (0.40 – 0.94). With regards to stages of LUAD tumours, higher *SCARF1* expression was strongly associated with better overall survival in early stage (Stage I) disease (HR = 0.27, 95 % CI = 0.12 – 0.59, *p* ≤ 0.001; Figure 4B), but held no prognostic value in more advanced (Stage II) cancers. High expression of *SCARF1* was also indicative of improved overall survival in non-smokers (HR = 0.37, 95 % CI = 0.13 – 1.03; Figure 4B), but exhibited no prognostic value in those patients with a history of smoking (Figure 4B). The level of *SCARF1* expression held no prognostic value in the presence or absence of lymph node metastasis in LUAD. In keeping with the fact that SCARF1 expression was not associated with tumour biology or outcome in any way, it was unsurprising that *SCARF1* showed no prognostic value in those patients with LUSC.

**Figure 4.**
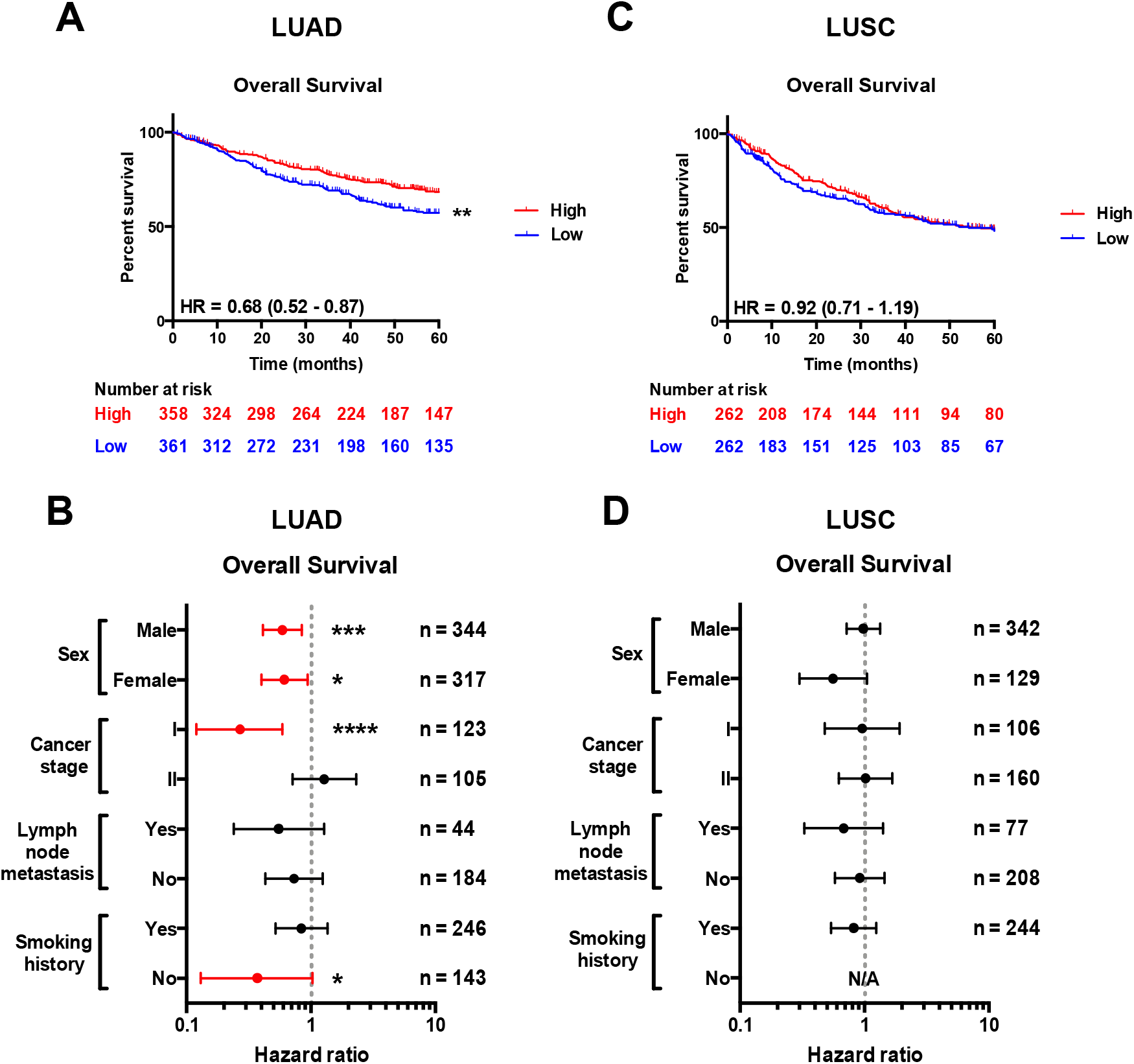
*SCARF1* expression is predictive of survival in LUAD, but not LUSC. (A) Overall survival in LUAD patients separated into two groups (‘High’ and ‘Low’ expression) via the median expression of *SCARF1.* ** indicates statistical significance where *p*□≤□0.01. HR = hazard ratio. (B) Forest plots of Overall survival in relation to various clinicopathological features of LUAD patients. Red plots highlight clinicopathological parameters in which statistical significance was achieved. *, *** and **** indicate statistical significance where *p*□≤□0.05, *p*□≤□0.005 or *p*□≤□0.001, respectively. (C) Overall survival in LUSC patients separated into two groups (‘High’ and ‘Low’ expression) via the median expression of *SCARF1.* (D) Forest plots of Overall survival in relation to various clinicopathological features of LUAD patients. Data in this Figure was generated with use of KM Plotter (http://kmplot.com/analysis/).

### SCARF1 *is expressed in tumour endothelial cells and correlates with CD4^+^ T cell infiltration in LUAD*

*SCARF1* exhibits a strong endothelial signature in HCC tumours [11] and, given that in homeostatic conditions, *SCARF1* is predominantly expressed in normal endothelial cell (NEC) populations within the human lung (Figure 1B), we next explored whether or not it was also expressed in tumour endothelial cells (TEC) of NSCLCs. With use of the publically-available single cell RNA sequencing data from the Lung Endothelial Cell taxonomy (LungECtax) database (https://endotheliomics.shinyapps.io/lungectax/), we were able to visualise the different endothelial cell populations present in normal human lung and NSCLC tissues. t-Distributed Stochastic Neighbour Embedding (tSNE) plots generated via the LungECtax software showed that there were 13 distinct endothelial cell subtypes present within human lung tissues (Figure 5A) and that certain populations were enriched in either normal or NSCLC tumour tissues (Figure 5B). The LungECTax software was subsequently used to explore the expression pattern of *SCARF1* across the various endothelial cell populations. *SCARF1* was ubiquitously expressed across throughout all 13 lung endothelial cell populations, showing comparable expression in both NEC- and TEC-enriched populations (Figure 5C). The expression of *SCARF1* in NSCLC tumour endothelia was highly consistent with our previous findings of its presence in HCC tumour endothelia.

**Figure 5.**
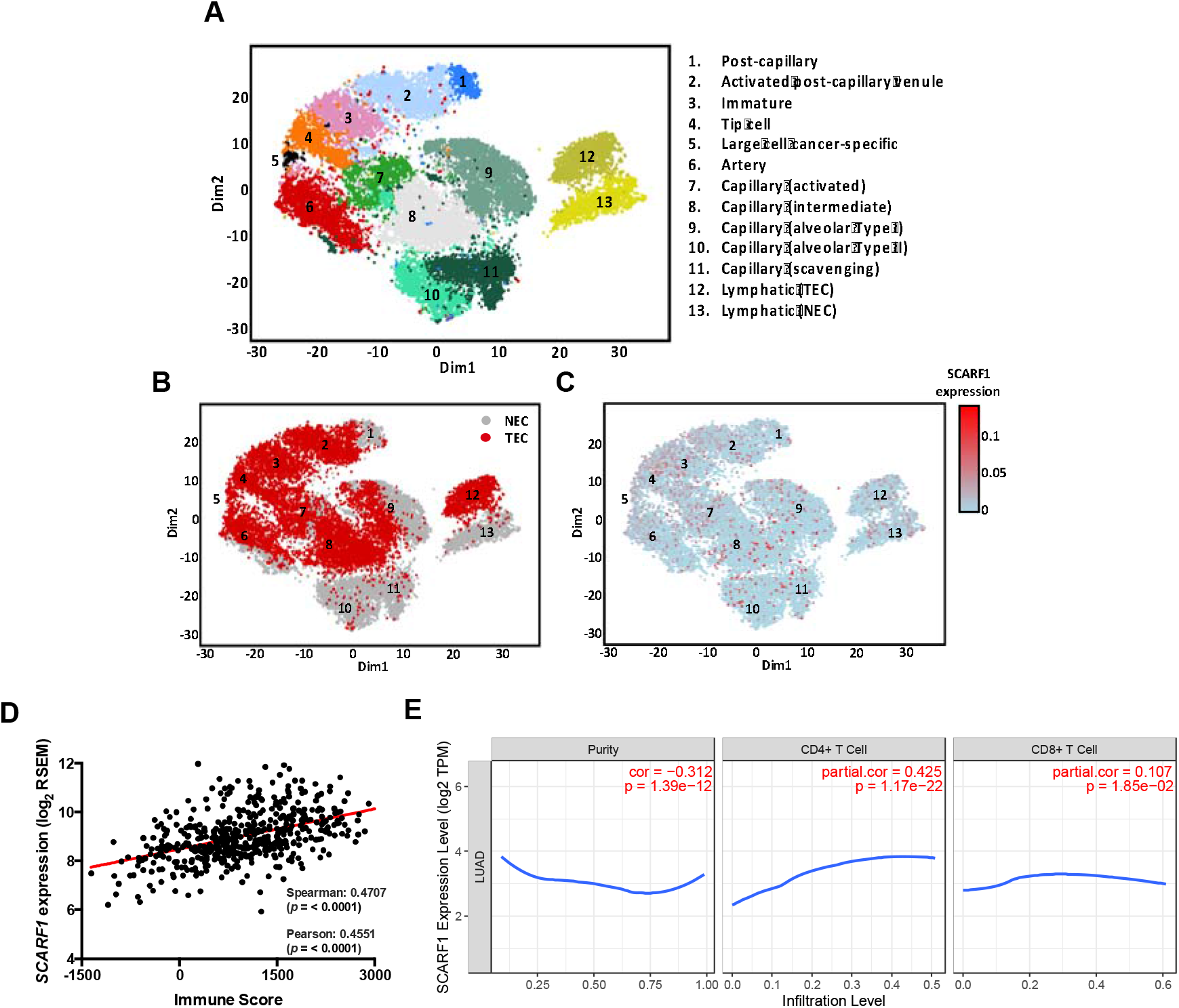
*SCARF1* is widely expressed in NSCLC tumour endothelial cell populations and correlates with CD4^+^ T cell infiltration in LUAD. (A) tSNE plot of endothelial cells subtypes present in normal and tumour human lung tissues. (B) tSNE plot of endothelial subtypes enriched in normal endothelial cells (NEC; grey) and tumour endothelial cells (TEC; red). (C) tSNE plot of *SCARF1* expression (red) in human lung endothelial cells subtypes. (D) Correlation of *SCARF1* expression with immune score in LUAD tumour tissues. (E) Correlation of *SCARF1* expression with the extent of CD4^+^ and CD8^+^ T cell infiltration in LUAD tumour tissues. tSNE plots in (A), (B) and (C) were generated by the publically-available LungECtax database (https://endotheliomics.shinyapps.io/lungectax/) accessed on 4^th^ June 2020. Data in (D) was generated via via Estimation of STromal and Immune cells in MAlignant Tumor tissues using Expression data (ESTIMATE; https://bioinformatics.mdanderson.org/estimate/; accessed 19th August 2020). Data in (E) was generated via the Tumor IMmune Estimation Resource (TIMER; https://cistrome.shinyapps.io/timer/; accessed 12th May 2020).

We have previously shown that *SCARF1* expression correlated with the level of CD4^+^ T cell infiltration in HCC tumours and hypothesised that it could function to specifically recruit proinflammatory subsets of CD4^+^ T cells [11]. Given the parallels of *SCARF1* in HCC and its expression in LUAD tumours described throughout the current study, we next aimed to investigate whether SCARF1 could play a role in the recruitment of tumour-infiltrating lymphocytes (TILs) to the LUAD tumour microenvironment. To explore this, we used a publically-available tool, Estimation of STromal and Immune cells in MAlignant Tumor tissues using Expression data (ESTIMATE; https://bioinformatics.mdanderson.org/estimate/) to correlate *SCARF1* expression with immune score, which is representative of immune infiltration in tumour tissues [34]. Using ESTIMATE, we demonstrated a moderate positive correlation between *SCARF1* expression and immune score in LUAD tumour tissues (Figure 5D). Next, to explore the relationship of *SCARF1* and TILs in more detail, we used Tumor IMmune Estimation Resource (TIMER; https://cistrome.shinyapps.io/timer/) to correlate *SCARF1* expression with the level of CD4^+^ T cell infiltration of LUAD tumours. Using TIMER, we first confirmed that *SCARF1* expression is absent from tumour cells, as indicated by a negative ‘purity’ correlation (−0.312, *p* = 1.39e^-12^; Figure 5E, *left panel).* Consistent with our previous findings in HCC, we also demonstrated a moderate positive correlation with CD4^+^ T cell infiltration (purity-corrected partial Spearman’s rho value = 0.425, *p* = 1.17e^-22^; Figure 5E, *middle panel*) in LUAD. To demonstrate its specificity for CD4^+^ T cells^23^, we also used TIMER to correlate *SCARF1* with CD8^+^ T cell infiltration in LUAD tumours and showed a much weaker correlation (purity-corrected partial Spearman’s rho value = 0.107, *p* = 1.85e^-02^; Figure 5E, *right panel).*

## Discussion

Genetic and molecular profiling of tumour tissues has aided the development of highly targeted and more effective treatments for lung cancers, significantly increasing patient survival time; however, lung cancer remains to be a major cause of mortality worldwide. Therefore, further investigation of the tumour microenvironment is warranted to identify novel biomarkers and therapies. Scavenger receptors have previously been show to bind a range of oncogenic ligands, such as heat shock proteins (HSPs) [35,36] and bacterial lipopolysaccharide (LPS) [37–39] and are known to be involved in the pathophysiology of a range of cancers [40]. Indeed, we have previously explored two such receptors, SCARF1 [11] and stabilin-1 [41], in the context of hepatocellular carcinoma (HCC) and hypothesised that they could play important roles in shaping the tumour microenvironment. Scavenger receptors in the lung are known to mediate key innate immune responses to a number of microbial infections [13,14] and environmental antigens [15–17]; however, the study of these receptors in the context of lung cancers remains relatively unexplored.

Recent work from our lab has focussed on the expression and function of a largely understudied scavenger receptor [23], SCARF1 (scavenger receptor class F, member 1), in human liver tissues [11,22]; however, others have also identified its presence in human lung tissues [18]. Here, we show that the lungs are a site of significant *SCARF1* expression (Figure 1A) and, like the liver [22], endothelial cells account for the majority of its expression (Figure 1B). In addition, consistent with our findings in gastrointestinal cancers, such as HCC and colonic adenocarcinoma [11], we showed here that *SCARF1* expression is significantly downregulated in two major histological subtypes of non-small cell lung cancer (NSCLC), lung adenocarcinoma (LUAD) and lung squamous cell carcionoma (LUSC) (Figures 1C-E). In addition, and once again consistent with our previous findings in HCC, we showed that higher intratumoural expression of *SCARF1* in LUAD tumour tissues was associated with less aggressive tumour biology (Figure 2) and better outcomes (Figure 3); however, in stark contrast, this association was not evident in LUSC tumours (Figures 2 & 3). Furthermore, from a prognostic perspective, higher *SCARF1* expression in LUAD tumours was highly indicative of better overall survival (Figure 4), but showed no prognostic value in LUSC tumours.

Endothelial-expressed scavenger receptors are known to possess a secondary function as atypical adhesion molecules in the leukocyte recruitment cascade [42] and we have recently described that SCARF1 is potentially involved in the recruitment of proinflammatory CD4^+^ T cells to the microenvironment of HCC tumours [11]. Thus, the disparities in the data between the two subtypes of NSCLC described in this study may be explained by the immune status of the two tumours types. LUAD tumours are known to possess a high clonal neoantigen burden [43], which is associated with an inflamed tumour microenvironment enriched in activated effector T cells [43] and higher expression of genes involved in antigen presentation, T cell migration and effector T cell function [43,44]. In stark contrast, LUSC tumours exhibit a low neoantigen burden [43] and are known to actively repress key genes in immunity, such as MHC molecules and chemokines, in early stage disease [44]. Ultimately, LUAD tumours may promote a more proinflammatory tumour microenvironment, whereas LUSC tumours are more likely to present as antiinflammatory to escape immune surveillance; therefore, we hypothesised that SCARF1 could function in LUAD tumours as it does in HCC. To explore this, we first needed to show that *SCARF1* was indeed present in tumour endothelial cells and we were able to do so via the publically-available Lung Endothelial Cell taxonomy (LungECtax) database (Figure 5A-C). Next, we showed a positive correlation between *SCARF1* expression and immune infiltration score (Figure 5D), and more specifically, the level of CD4^+^ T cell infiltration in LUAD tumours (Figure 5E, *middle panel*). Finally, we corroborated our previous findings of SCARF1’s specificity for CD4^+^ T cells [22], by demonstrating a negligible correlation between LUAD-expressed *SCARF1* and CD8^+^ T cell infiltration (Figure 5E, *right panel*). This is particularly relevant to the context of LUAD, as previous studies have shown that an increased prevalence of tumour-infiltrating lymphocytes is a prognostic factor in NSCLCs [45,46]. Therefore, increasing the inflammatory status via a SCARF1-mediated pathway could have a significant impact on patient outcome.

Although this study represents the first detailed investigation of *SCARF1* in human lung tissues and their associated malignancies, this work is still very preliminary as it has only been undertaken with publically-available RNA sequencing data. Therefore, future work should initially focus on corroborating these data with protein expression experiments on *ex vivo* sections of human lung tissue. In addition, flow-based adhesion assays with primary lung endothelial cells, as we have previously performed with liver endothelial cells [47], would confirm functionality of lung-expressed SCARF1. Should this be consistent with the data presented in the current study, then further *in vivo* work would then be required with SCARF1 knockout models to fully confirm its contribution to the NSCLC immune microenvironment. Taken all together, our data presented here describes a potential beneficial role for SCARF1 in LUAD tumours. The current study, along with previous data, further emphasises a role for SCARF1 in shaping the tumour microenvironment and is highly suggestive that SCARF1 present novel therapeutic target in the treatment of immunologically active cancers.

## Author Contributions

Conceptualization, DAP; Data curation, DAP; Formal analysis, DAP; Funding acquisition, DAP and SS; Investigation, DAP; Methodology, DAP; Writing – original draft, DAP; Writing - review & editing, DAP and SS.

## Acknowledgments

This paper presents independent research supported by the Birmingham NIHR Liver Biomedical Research Unit based at the University Hospitals Birmingham NHS Foundation Trust and the University of Birmingham. The views expressed are those of the authors and not necessarily those of the NHS, the NIHR or the Department of Health.

